# Mixing crop residues induces a synergistic effect on microbial biomass and an additive effect on soil organic matter priming

**DOI:** 10.1101/2021.05.11.443543

**Authors:** Xin Shu, Yiran Zou, Liz J. Shaw, Lindsay Todman, Mark Tibbett, Tom Sizmur

**Affiliations:** Department of Geography and Environmental Science, University of Reading, UK; Department of Sustainable Land Management and Soil Research Centre, School of Agriculture, Policy and Development, University of Reading, UK

**Author notes:** Corresponding author –.

**Keywords:** diverse, mixture, synergistic, ^13^C-PLFA, crop residues

## Abstract

Applying crop residues is a widely used strategy to increase soil organic matter (SOM) in arable soils because of its recorded effects on increasing microbial biomass and consequently necromass. However, fresh residue inputs could also “prime” the decomposition of native SOM, resulting in accelerated SOM depletion and greenhouse gas (GHG) emission. Increasing the botanical diversity of the crops grown in arable systems has been promoted to increase the delivery of multiple ecological functions, including increasing soil microbial biomass and SOM. Whether mixtures of fresh residues from different crops grown in polyculture contribute to soil carbon (C) pools to a greater extent than would be expected from applying individual residues (i.e., the mixture produces a non-additive synergistic effect) has not been systematically tested and is currently unknown. In this study, we used ^13^C isotope labelled cover crop residues (i.e., buckwheat, clover, radish, and sunflower) to track the fate of plant residue-derived C and C derived from the priming of SOM in treatments comprising a quaternary mixture of the residues and the average effect of the four individual residues one day after residue incorporation in a laboratory microcosm experiment. Our results indicate that, despite all treatments receiving the same amount of plant residue-derived C (1 mg^-1^ C g soil), the total microbial biomass in the treatment receiving the residue mixture was significantly greater, by 26% (3.69 µg^-1^ C g), than the average microbial biomass observed in treatments receiving the four individual components of the mixture one day after applying crop residues. The greater microbial biomass C in the quaternary mixture, compared to average of the individual residue treatments, that can be attributed directly to the plant residue applied was also significantly greater, by 132% (3.61 µg^-1^ C g). However, there was no evidence that the mixture resulted in any more priming of native SOM than average priming observed in the individual residue treatments. The soil microbial community structure, assessed using phospholipid fatty acid (PLFA) analysis, was significantly (*P* < 0.001) different in the soil receiving the residue mixture, compared to the average structures of the communities in soil receiving four individual residues. Differences in the biomass of fungi, general bacteria, and Gram-positive bacteria were responsible for the observed synergistic effect of crop residue mixtures on total microbial biomass and residue-derived microbial biomass, especially biomarkers 16:0, 18:2ω6 and 18:3ω3. Our study demonstrates that applying a mixture of crop residues increases soil microbial biomass to a greater extent than would be expected from applying individual residues and that this occurs either due to faster decomposition of the crop residues or greater carbon use efficiency (CUE), rather than priming the decomposition of native SOM. Therefore, growing crop polycultures (e.g., cover crop mixtures) and incorporating mixtures of the resulting crop residues into the soil could be an effective method to increase microbial biomass and ultimately C stocks in arable soils.

## 1. Introduction

Soil organic carbon (SOC) plays a critical role in global carbon (C) dynamics in the earth system and is a major property influencing soil functions and health. Applying crop residues to soils is a common strategy used in agroecosystems to enhance SOC stocks (Chapman and Newman, 2010; Chen et al., 2015). When microorganisms decompose plant litter and use the C for metabolism, they catabolise a portion of this C, which is usually released as carbon dioxide (CO_2_), and simultaneously assimilate and anabolise a portion of C into their biomass. After their death, microbial necromass returns to soil and contributes to SOC storage (Liang et al., 2020). However, applying fresh crop residues could also stimulate microbial decomposition of native SOM via co-metabolism or mining for nutrients, resulting in a priming effect (Kuzyakov, 2010; Wang et al., 2015). A recent meta-analysis including 2048 individual experimental comparisons from 94 laboratory incubation studies revealed that the addition of exogenous organic C significantly enhanced native SOC decomposition, by 47.5%, in terrestrial ecosystems, with the highest priming effect observed in arable soils (60.9%) (Sun et al., 2019). Therefore, using crop residues to maximize SOC stocks requires a consideration of the impact of amendments on SOC priming.

Previous studies have investigated the C mass balance after the application of plant residues of a single plant species (Rubino et al., 2010; Shahbaz et al., 2018). However, crop residues returned to soils under arable land management practices such as intercropping, rotations, and cover crops include the residues of more than one plant species. Based on studies examining decomposition dynamics in soils receiving mixed species residues, it cannot necessarily be assumed that the application of a crop residue mixture will have the same impact on C dynamics as predicted from observations made on the impact of the individual residues (Gartner and Cardon, 2004; Porre et al., 2020). If the behaviour or effect of a residue mixture can be predicted from the behaviour of the individual residues, this is classified as an additive effect (Redin et al., 2014). By contrast, a mixture could also deliver an antagonistic non-additive effect (i.e. the mixture’s effect is less than the average of individual species) or a synergistic non-additive effect (i.e. the mixture’s effect is greater than the average of individual species), which suggests there are interactions, via microbial decomposers, among the constituents of the mixture (Redin et al., 2014).

A majority of the previous studies have explored the effects of mixtures on litter decomposition and associated C cycling and nutrient release (e.g. nitrogen mineralization and immobilization) by focusing on leaf litter decomposition in forest ecosystems (Castro-Díez et al., 2019; Gartner and Cardon, 2004; Mao et al., 2017). Even the most recent meta-analysis on litter mixtures only focused on litter mass loss due to decomposition (Porre et al., 2020). To better understand the mechanisms of plant species diversification on soil C dynamics, we need to fill the knowledge gap regarding mixture effects on the microbial fate of C supplied by individual components of the mixture and the potential for interactions of residue C with older soil organic matter (SOM) via priming effects.

The mechanisms responsible for non-additive effects in residue mixture decomposition dynamics are not fully understood and might depend strongly on the context within which the study was conducted (Porre et al., 2020). To explain non-additive effects, processes relating to nutrient transfer between nutrient-rich (low C:N) and nutrient poor (high C:N) litters, transfer of inhibitory compounds from one species’ litter to another, or physical (water retention) effects have been frequently mentioned (Porre et al., 2020). In addition, mixing chemically contrasting crops may provide a greater number of niches for microorganisms to exploit, which allows functionally dissimilar microbial communities to coexist, and thus result in a greater microbial diversity and biomass than might be expected from the average of the individual communities that are supported by monocultures (Chapman and Newman, 2010). Although greater diversity not always linked to greater biomass, a recent meta-analysis found that in temperate regions, where soil is not rich in C, greater microbial diversity is usually associated with greater microbial biomass due to facilitation and niche partitioning by supporting the co-existence of multiple microbial species (Bastida et al., 2021). An increased microbial diversity could increase the probability of including taxa particularly influential in extracellular enzyme production for nutrient mining and SOM degradation through metabolic and co-metabolic processes. Furthermore, an increased microbial biomass could also result in stronger SOM mineralization as a consequence of increased microbial metabolic activity (Bastida et al., 2021). Thus, mixtures of crop residues could enhance the extent to which soil microbial communities mine nutrients for SOM and lead to a greater priming effect. By contrast, some studies showed that mixtures of diverse plant residues have a greater opportunity to provide preferable growth substrates for microbes (e.g., if the average C/N ratio of residues is close to 24), thereby offsetting the extent to which SOM decomposition is primed and decreasing microbial anabolism of primed SOM (Xiao et al., 2015). These two mechanisms (i.e., increasing or offsetting priming effect) could occur simultaneously and combine to determine the overall magnitude of the priming effect.

In this study, we investigated the residues of four functionally dissimilar crops from four different plant families (i.e., buckwheat, clover, radish, and sunflower) which are widely grown in mixtures as cover crops in agricultural systems. We established a microcosm experiment comprising treatments receiving either mixtures or individual (non-mixture) ^13^C labelled cover crop residues which provided the same amount of residue-derived C (1 mg^-1^ C g soil). Soil phospholipid fatty acid (PLFA) analysis was undertaken one day after incorporating crop residues to quantify the biomass of key soil microbial groups. Gas chromatography-combustion-stable isotope mass spectrometry (GC-C-IRMS) was used to identify the microbial groups that had incorporated residue-derived C and, by mass balance, quantify the amount of primed SOM-derived C which was incorporated into the microbial biomass. The difference between the mixture and the average of four non-mixtures enabled us to determine whether the mixture delivered either a synergistic (mixture > average), an antagonistic (mixture < average), or an additive (mixture = average) effect.

We assumed that microbes have no preference for ^13^C over ^12^C. Because a mixture of crop residues may increase the niche breadth and provide a more diverse supply of nutrients, thereby creating conditions enhancing the growth and facilitation of multiple co-exist microorganisms. Thus, we hypothesised that the mixture could result in a synergistic effect on total microbial biomass. Compared to individual species, the mixture has higher probability to provide preferable growth substrates which could be more easily assimilated into microbial biomass, and thus we would expect a synergistic effect of the mixture on the microbial biomass derived from plant residues. Given that the cover crop species tested had divergent C:N ratios (ranging between 10 and 32 and spanning the threshold C:N (≈24) for net N mineralization-immobilization (Norton and Schimel, 2011), we hypothesised that adding residues in a mixture (average C/N = 17) would decrease the requirement for microorganisms to prime native SOM to scavenge for N and therefore induce an antagonistic effect on the microbial biomass C derived from primed SOM.

### 2. Materials and methods

### 2.1. Soil samples and crop residues

A silty loam Luvisol (World Reference Base classification); pH (H_2_O) 6.3, 22.32 g C kg^-1^, 2.24 g N kg^-1^, 0.90 mg NH_4_ ^+^-N kg^-1^, 2.75 mg NO_3_ -N kg^-1^ was collected from an arable field on the University of Reading’s research farm at Sonning, Reading, UK (51.481152, -0.902188) in August 2019 after harvesting spring barley (*Hordeum vulgare*). Seven surface soil samples (0-20 cm depth) were randomly sampled and mixed thoroughly to create one homogenous sample, approximately 20 kg in weight.

Four cover crops, buckwheat (*Fagopyrum esculentum*), berseem clover (*Trifolium alexandrinum*), oil radish (*Raphanus raphanistrum*), and sunflower (*Helianthus annuus*), were continuously and uniformly labelled with ^13^CO_2_ in growth cambers by IsoLife (Wageningen, Netherlands). Buckwheat and clover were harvested 5 weeks after sowing, while radish and sunflower were harvested after 4 weeks. The ^13^C atom percent of the resulting aboveground biomass was 6.7%, 7.8%, 7.8%, and 8.0% for buckwheat, clover, radish, and sunflower, respectively. Corresponding unlabelled crops were grown under the same conditions in growth chambers by IsoLife (Wageningen, Netherlands), and harvested at the same time. After harvesting, the aboveground residues of both ^13^C labelled and unlabelled crops were dried at 70 °C and milled to pass through 0.05 mm mesh. The chemical composition of ^13^C labelled and unlabelled residues is provided in Table S1.

### 2.2. Experimental design

Soil was sieved to pass a 4 mm mesh and then pre-incubated for 7 days at 26 °C with a soil water content of 60% of the water holding capacity (0.22 g g^-1^). As indicated in Table 1, the treatments consisted of pure unlabelled residues, labelled non-mixture residues, quaternary mixtures of residues which contained one labelled species and three unlabelled species, and a control without any crop residue additions. For each treatment receiving residues, four replicate microcosms were established by mixing 150 g of fresh soil (equivalent to 122.95 g dry soil) thoroughly with a mass of dry residues to ensure C was added to each microcosm at a rate of 1 mg C g^-1^ soil. In the pure treatments, all the added C was from the same unlabelled residue sample. In the non-mixture treatments, 25% of added C was from the ^13^C labelled residue and 75% of added C was from the unlabelled residue of the same crop species. In the mixture treatments, 25% of added C was from a ^13^C labelled crop and 75% of added C comprised unlabelled residues from the other three crop species.

**Table 1.** Experimental design and the C/N ratio of added crop residues in each treatment.

| Treatment | Plant | Abbreviation | Description | Added <sup>13</sup>C amount<br>(mg C g<sup>-1</sup> soil) | C/N ratio |
| --- | --- | --- | --- | --- | --- |
| Non-mix | Buckwheat | NB | 25% labelled + 75% unlabelled buckwheat | 0.0168 | 10 |
| Non-mix | Clover | NC | 25% labelled + 75% unlabelled clover | 0.0195 | 32 |
| Non-mix | Radish | NR | 25% labelled + 75% unlabelled radish | 0.0195 | 18 |
| Non-mix | Sunflower | NS | 25% labelled + 75% unlabelled sunflower | 0.0200 | 19 |
| Mix | Buckwheat | MB | 25% labelled buckwheat, 75% of unlabelled residues (clover, radish, sunflower) | 0.0168 | 17 |
| Mix | Clover | MC | 25% labelled clover, 75% unlabelled residues (buckwheat, radish, sunflower) | 0.0195 | 17 |
| Mix | Radish | MR | 25% labelled radish, 75% unlabelled residues (buckwheat, clover, sunflower) | 0.0195 | 17 |
| Mix | Sunflower | MS | 25% labelled sunflower, 75% unlabelled residues (buckwheat, clover, radish) | 0.0200 | 17 |
| Pure | Buckwheat | PB | %100 unlabelled buckwheat | \ | 9 |
| Pure | Clover | PC | %100 unlabelled clover | \ | 30 |
| Pure | Radish | PR | %100 unlabelled radish | \ | 21 |
| Pure | Sunflower | PS | %100 unlabelled sunflower | \ | 22 |
| Soil |  |  | Soil only without any plant residue addition | \ | \ |

*Note: 25% and 75% refers to proportion of total added C. The quantity of C added was the same across all treatments apart from the soil treatment, which* *was 1 mg C g<sup>-1</sup> soil. The C/N ratio in the mixture treatments were the average of four types of mixture treatments.*

For the measurement of soil respiration, a 100 g subsample was transported from each microcosm to a bulk density ring (98 cm^3^), stored in a gas-tight plastic jar (365 cm^3^), and kept open during incubation. The rest of sample was kept in an opened plastic bag for the measurement of PLFA. Both measurements of PLFA and soil respiration were taken one day after applying cover crop residues.

To measure soil respiration, jars were sealed with a Suba-Seal^®^ Septa for 1 h and a 16 ml headspace gas sample was taken from each jar using a syringe and hypodermic needle, transferred into pre-evacuated vials, and analysed with gas chromatography (Agilent 7890B, UK) (Adekanmbi et al., 2020). The universal gas law was used to determine the amount of CO_2_ (ng g^-1^ soil h^-1^) emitted from each jar.

### 2.3. Phospholipid-derived fatty acids (PLFA) extraction

One day after incubation with soil and crop residues, a 10 g aliquot of soil was sampled from each replicate plastic bag and freeze-dried for downstream analysis of PLFA. PLFA was extracted following the method described by Sizmur et al. (2011). Briefly, 4 g of freeze-dried soil was extracted with 7.8 ml of Bligh and Dyer extractant containing chloroform: methanol: citrate buffer (1:2:0.8 v/v/v). The extracted phospholipids were methanolized as fatty-acid methyl esters and dissolved in hexane for analysis by gas chromatography (GC).

### 2.4. Gas chromatography (GC)

PLFA methyl esters were analysed using an Agilent Technologies 6890N gas chromatography equipped with a Supercowax 10 capillary GC column (60 m × 0.25 mm i.d. × 0.25 μm film thickness) and a Flame Ionisation Detector (FID). Helium was the carrier gas. The temperature programme was 1-minute isothermal at 60 °C, followed by a ramp to 145 °C at 25 °C per minute, followed by an increase to 250 °C at 2.5 °C per minute and then held isothermally at 310 °C for 10 minutes. Data were processed using GC ChemStation (Agilent Technologies). Peaks were identified using a bacterial fatty acid methyl esters (BAME) mix (Sigma Aldrich, UK) and quantified using a 37-component fatty acid methyl esters (FAME) mix (Sigma Aldrich, UK). The biomass of each group of microorganisms was determined using the combined mass of fatty acids to which the group is attributed in Table S2.

### 2.5. Gas chromatography-combustion-isotope ratio mass spectrometry (GC-C-IRMS)

GC-C-IRMS analysis were performed by injecting a 1 µl sample of fatty-acid methyl esters into an Agilent 7890N GC, upstream of a DELTA V™ Isotope Ratio Mass Spectrometer (electron ionization, 100 eV, 1 mA electron energy, 3 F cup collectors m/z 44, 45 and 46, CuO/Pt Thermofisher GC IsoLink interface maintained at 1000°C). A Nafion membrane was employed to prevent water from reaching the ion source. GC conditions were the same as that described above. Samples were calibrated against reference CO_2_ of known isotopic composition, which was introduced directly into the source five times at the beginning and end of every run. Data was processed in the Isodat Gas Isotope Ratio MS Software (ThermoFisher Scientific) to generate δ^13^C values representing the ratio of ^13^C/^12^C in fatty-acid methyl esters, relative to the ^13^C/^12^C ratio of the international Pee Dee Belemnite (PDB) standard (0.01118).

δ^13^C values obtained for methylated compounds were corrected for the addition of derivative C using the mass balance equation following equation 1 to 3 (Zhang et al., 2019):

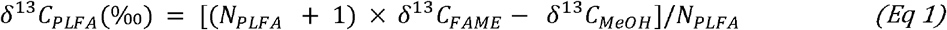

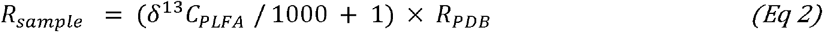

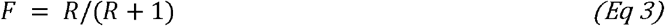

where δ^13^C_PLFA_, δ^13^C_FAME_, and δ^13^C_MeOH_, were the δ^13^C value of PLFA, FAME and methanol (δ^13^C_MeOH_ was assumed to equal zero), respectively. N_PLFA_ was the number of C atoms in this fatty acid to which the fatty acid methyl ester corresponds. R_sample_ and R_PDB_ were the ratios of ^13^C/^12^C in sample and in PDB. F is the fractional isotopic abundance representing the concentration of ^13^C as a proportion of the total concentration of C in the PLFA.

Equation 4 was used to calculate how much ^13^C (µg ^13^C g^-1^ soil) was incorporated into PLFA biomass in the non-mixture treatments:

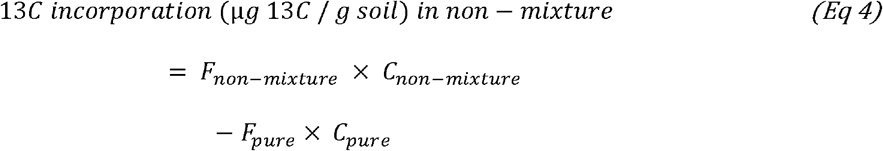

F_non-mixture_ and F_pure_ were the fractional abundances of ^13^C in a non-mixture treatment its corresponding pure unlabelled treatment, respectively. C_non-mixture_ and C_pure_ were the concentration of PLFAs (µg C g^-1^ soil) in the non-mixture and its corresponding pure unlabelled treatment, which were the results of GC-FID analysis. For example, ^13^C incorporation in the non-mixture of buckwheat (NB) equals to *F*_*NB*_ × *C*_*NB*_ −*F*_*PB*_ ×*C*_*PB*_

For each mixture treatment, there was 75% of added C from three different unlabelled residues (25% from each residue). The resulting ^13^C incorporation (µg ^13^C g^-1^ soil) of these unlabelled residues was accounted for using equation 5.

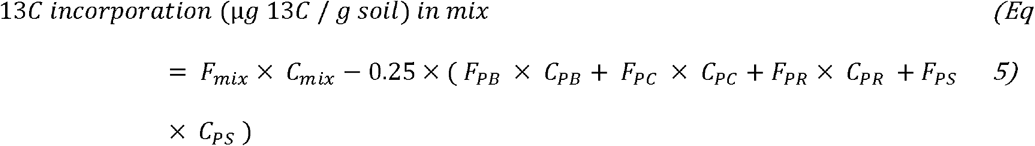

where F_mix_ was the fractional abundances of ^13^C in mixture treatment. F_PB_, F_PC_, F _PR_, F _PS_, were the fractional abundances of ^13^C in pure unlabelled buckwheat, clover, radish, and sunflower treatment, respectively. C_mix_, C_PB_, C_PC_, C_PR_, C_PS_, were the concentration of PLFAs (µg C g^-1^ soil) in the treatments of mixture, pure buckwheat, pure clover, pure radish, and pure sunflower, which were analysed by GC-FID.

The proportion (unitless) of ^13^C incorporated from total amount of added ^13^C was calculated using equations 6 and 7:

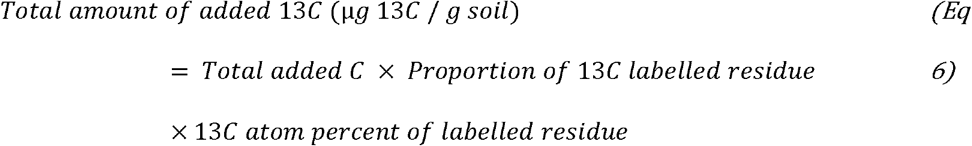

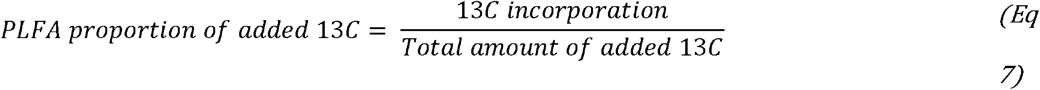

where total added C was 1000 µg C g^-1^ soil, the proportion of ^13^C labelled residue was 0.25. The ^13^C atom percent was 6.7%, 7.8%, 7.8% and 8.0% for buckwheat, clover, radish, and sunflower residues, respectively. This brings the total amount of added ^13^C to 16.75, 19.50, 19.50, and 20.00 µg ^13^C g^-1^ soil, for treatments which received 25% of their added C from ^13^C labelled buckwheat, radish, clover, and sunflower.

^13^C is a tracer to track the amount of C derived from crop residues and incorporated into PLFA biomass. Assuming microorganisms have no preference for ^12^C over ^13^C, the proportion of PLFA incorporated ^13^C from added ^13^C should also be the same to the proportion of PLFA incorporated ^12^C from total added ^12^C. Following this logic, in all the non-mixture treatments, the amount of C incorporated into PLFA derived from added crop residue C was calculated using equation 8:

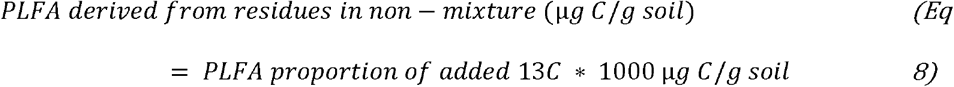

where 1000 µg C g^-1^ soil was the total added amount of C (equal to 1 mg C g^-1^ soil).

In the mixture treatment, the ^13^C labelled residue contributed 25% of the added C (i.e., 250 µg C g^-1^ soil). The proportion of ^13^C in each type of mixture (calculated by equation 7) was multiplied by 250 (µg C g^-1^ soil) to get the amount of PLFA derived from this ^13^C labelled crop. We then added up the amount of PLFA derived from each type of ^13^C labelled crop to form the PLFA which was derived from the mixture of crop residues (equation 9).

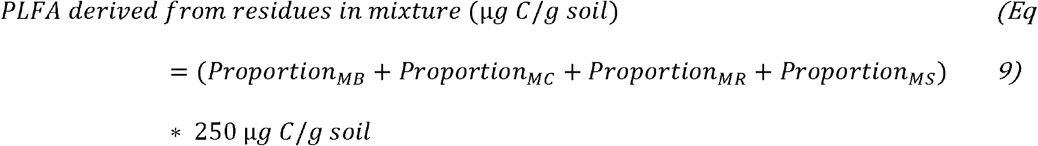

where proportion_MB_, proportion_MC_, proportion_MR_, and proportion_MS_ represent the proportion (unitless) of ^13^C incorporated from added ^13^C in four types of mixture treatments where each type was ^13^C-labelled by buckwheat, clover, radish, and sunflower, separately.

The PLFA derived from primed SOM was calculated as the difference between the total PLFA (measured using GC-FID), PLFA derived from residues, and PLFA in the control soil without any residue addition using equation 10.

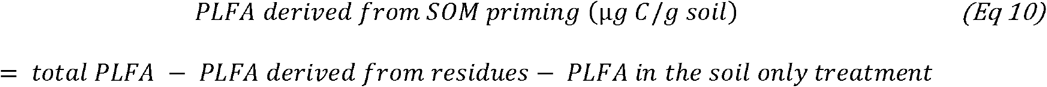

where total PLFA and PLFA in the soil only treatment were analysed by GC-FID and converted to µg C g^-1^ soil.

For mixture treatment, because there were only four samples have crop residue-derived PLFA, to calculate SOM-derived PLFA, we created a general mixture of which total PLFA was the average of total PLFA in four different types of mixture in the same replicate as calculated by equation 11.

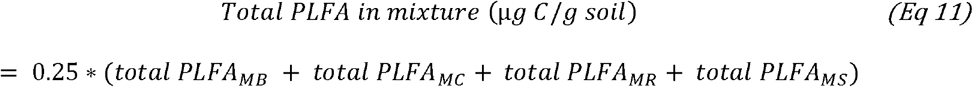

Therefore, there were four replicates for total PLFA, residue derived PLFA, and SOM derived PLFA in the general mixture.

### 2.6. Data analysis

All the statistics were conducted in R (version 3.5.2) (R Core Team, 2018) except for the analysis of similarity (ANOSIM) which was conducted in Primer 7 (Primer-e, Auckland, New Zealand). We used nested analysis of variance (ANOVA) to compare the effect from the quaternary mixtures with the effect from the average of the four non-mixture treatments on total PLFA biomass, the PLFA biomass incorporated from crop residues, and the PLFA biomass incorporated from primed SOM.

To compare the difference in microbial community structure between the mixture and the average of four individuals, PLFA data measured by GC-FID was “Hellinger” transformed in all treatments. Non-metric multidimensional scaling (NMDS) on Bray-Curtis distance of the transformed data was performed using the “vegan” package in R (Oksanen et al., 2019). Bray-Curtis distance similarity matrices were analysed by a one-way analysis of similarity (ANOSIM) using Primer 7 to test if the differences between the mixture and the average of four non-mixture treatments were significant.

## 3. Results

### 3.1 Crop residues increased soil microbial biomass and altered community structure

The incorporation of cover crop residues significantly (*P* < 0.001) shifted the microbial community structure away from the unamended control soil (Figure S1). The incorporation of residues from different crop species lead to significantly (*P* < 0.05) different community structures (Figure S1). The biomass of general bacteria, Gram-positive bacteria, and fungi in the unamended control soil was 1.80, 2.20, and 2.36 µg C g^-1^, respectively, which was greater than the biomass of Gram-negative bacteria and protozoa (Table S3). Despite the same rate of C addition applied across all the treatments, total PLFA biomass differed between residue amendment treatments; ranging from 11.66 to 18.93 µg C g^-1^, which was significantly (*P* < 0.05) greater than that in the control soil (Table S3).

### 3.2 Mixing crop residues increased total PLFA biomass and altered microbial community structure

Total PLFA biomass was 17.74 and 14.05 µg C g^-1^ for the mixture and the average of four non-mixture treatments, respectively (Figure 1). The soils in the mixture treatments had a significantly (*P* < 0.05) greater total microbial biomass, by 26% (3.69 µg C g^-1^), compared to the soils in the non-mixture treatments (Figure 1 and Table 2).

**Figure 1.**
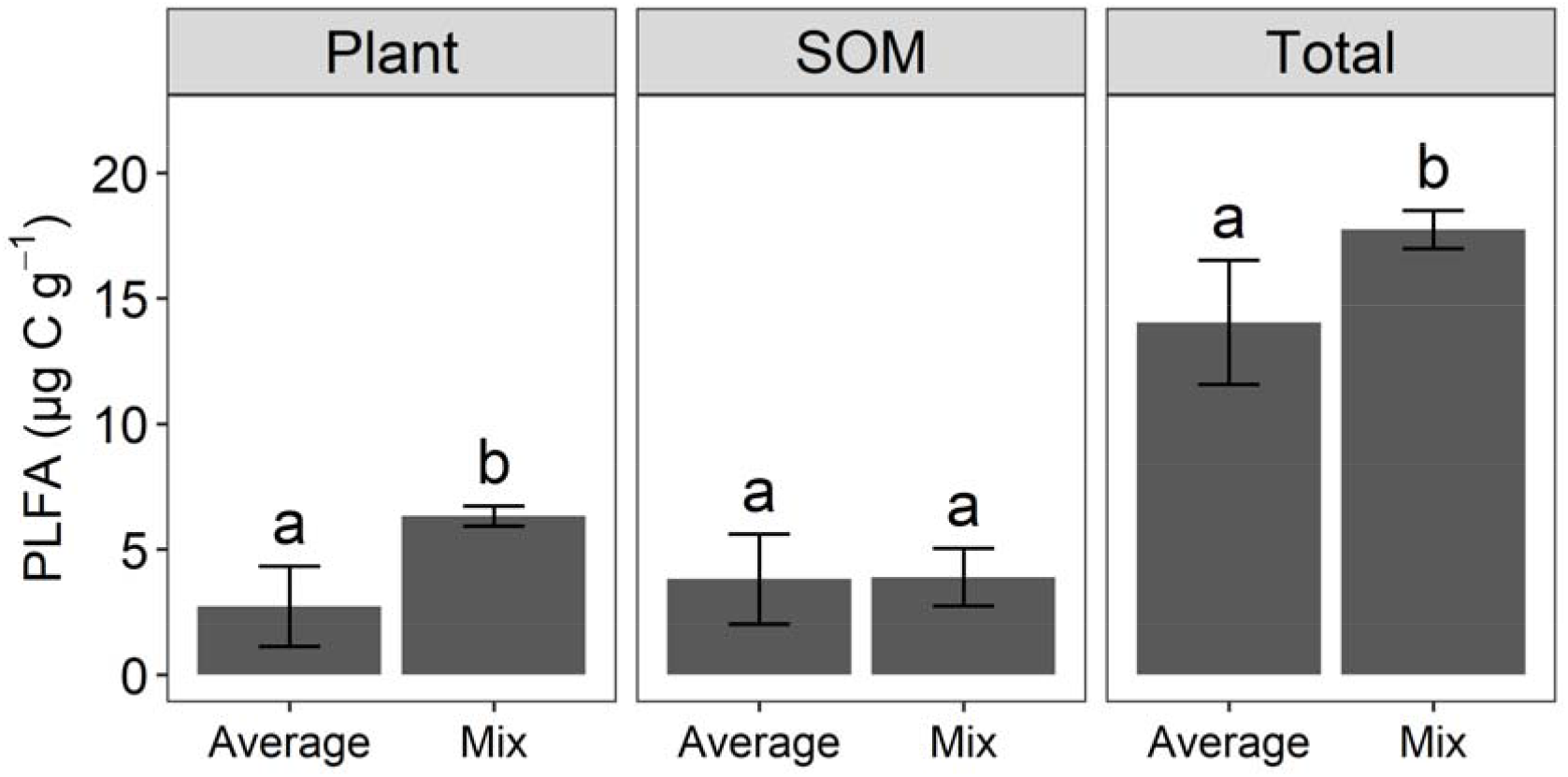
Total PLFA biomass (Total), PLFA biomass incorporated from crop residues (Plant), and the PLFA biomass incorporated from soil organic matter (SOM) in the mixture and the average of four non-mixture treatments. Error bars are standard deviations. Different letters within the same panel indicate a significant difference between treatment means at P <0.05.

**Table 2.** NEST ANOVA (treatment nested with crop species) to compare the effects of mixture and the species of plant on total PLFA, crop residue-derived PLFA, and SOM (soil organic matter)-derived PLFA at different microbial groups. MIX has two levels (mix or non-mix), crop species have five levels (radish, clover, buckwheat, sunflower, and mixture). DF is the degree of freedom. Values are F value. *, **, and *** represent significant level at *P* < 0.05, 0.01, and 0.001. G+ and G-bacteria are Gram-positive and Gram-negative bacteria, respectively.

| **Total PLFA** |  |  |  |  |  |  |  |  |  |
| --- | --- | --- | --- | --- | --- | --- | --- | --- | --- |
|  | DF | Sum of PLFA | General bacteria | G+ bacteria | G- bacteria | Fungi | Protozoa | G+/G- | Fungi/Bacteria |
| Treatment | 1 | 15.16** | 9.16** | 12.54** | 0.07 | 9.66** | 0.12 | 0.17 | 1.80 |
| Treatment:Species | 3 | 5.78** | 2.13 | 9.55*** | 4.32* | 2.69 | 0.95 | 3.46* | 4.01* |
| **Crop residue- derived PLFA** |  |  |  |  |  |  |  |  |  |
| Treatment | 1 | 20.62*** | 16.93*** | 4.64* | 0.19 | 11.43** | 0.22 | 0.40 | 0.02 |
| Treatment:Species | 3 | 1.36 | 0.12 | 1.34 | 2.66 | 1.41 | 0.85 | 0.71 | 0.4 |
| **SOM- derived PLFA** |  |  |  |  |  |  |  |  |  |
| Treatment | 1 | 0.01 | 0.07 | 1.33 | 0.00 | 0.03 | 0.01 | 0.44 | 0.53 |
| Treatment:Species | 3 | 2.72 | 2.14 | 3.97* | 3.53* | 1.48 | 0.81 | 1.08 | 2.63 |

The mixture treatment resulted in a significantly (*P* < 0.001) greater general bacteria, Gram-positive bacteria, and fungi biomass by 31% (1.07 µg C g^-1^), 18% (0.62 µg C g^-1^), and 38% (1.90 µg C g^-1^), respectively, than the average of the non-mixture treatments (Figure 2). In particular, biomarkers i15:0, 16:0, 18:1ω9, 18:2ω6 and 18:3ω3 were significantly (*P* < 0.05) more abundant in the mixture treatment than the average of four non-mixture treatments (Figure 3). The ratios of fungi to bacteria in the mixture and the average of the four non-mixture treatments were 0.64 and 0.57, which was not significantly different (Figure S2). Similarly, there was no significant difference in the ratio of Gram-positive to Gram-negative bacteria between the mixture and the average of four non-mixture treatments (Figure S3).

**Figure 2.**
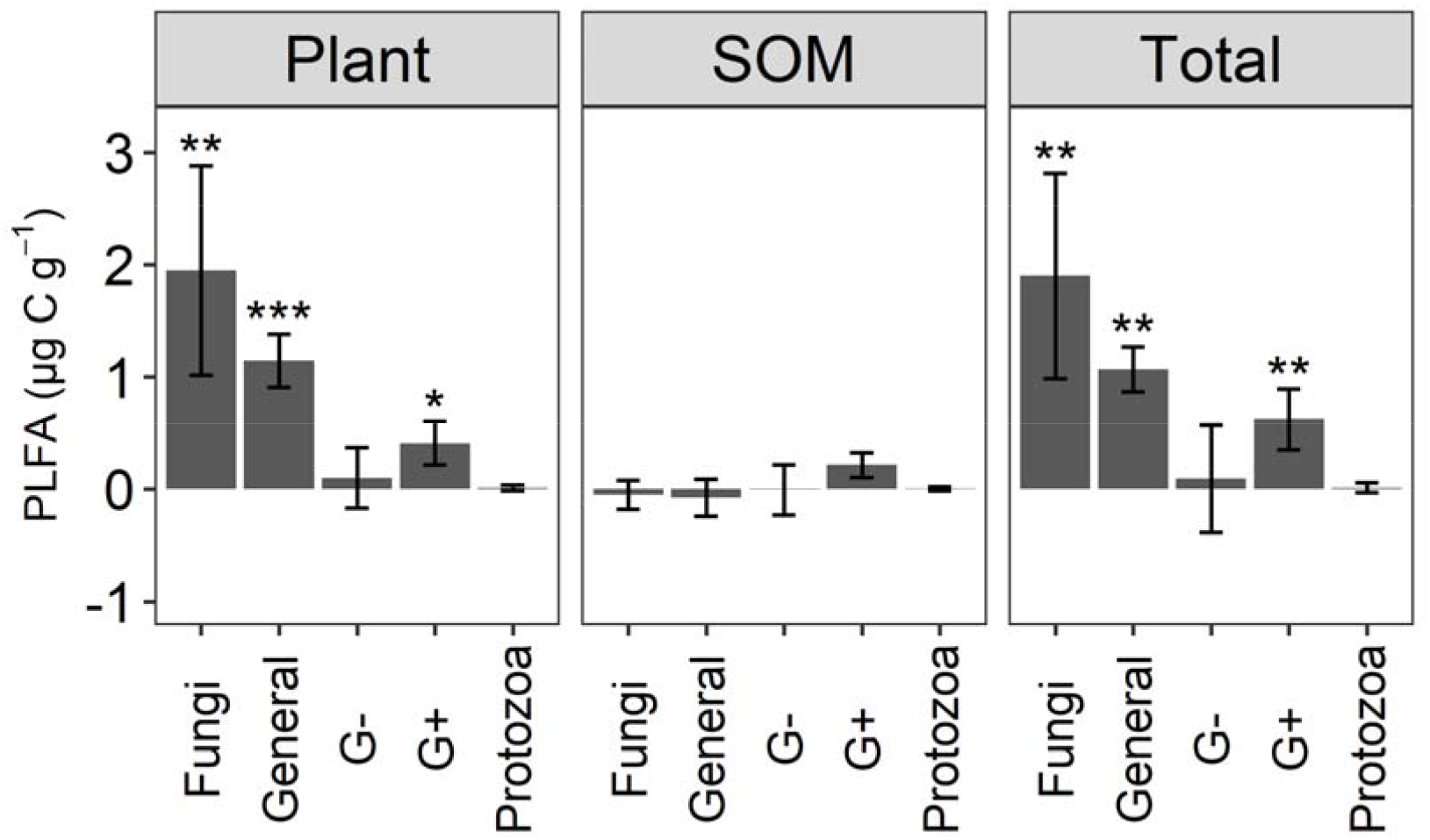
Differences in PLFA biomass of key microbial groups between the mixture and the average of four non-mixture treatments. Positive value means mixture treatment has greater biomass than the average of non-mixture treatments. The three panels represent the crop derived PLFA biomass (Plant), the primed SOM derived PLFA biomass (SOM), and the total PLFA biomass (Total). Error bars are standard deviations. General, G+, and G-represent general bacteria, Gram-positive and Gram-negative bacteria, respectively. *, **, or *** means significantly different from zero at the level P < 0.05, 0.01 or 0.001.

**Figure 3.**
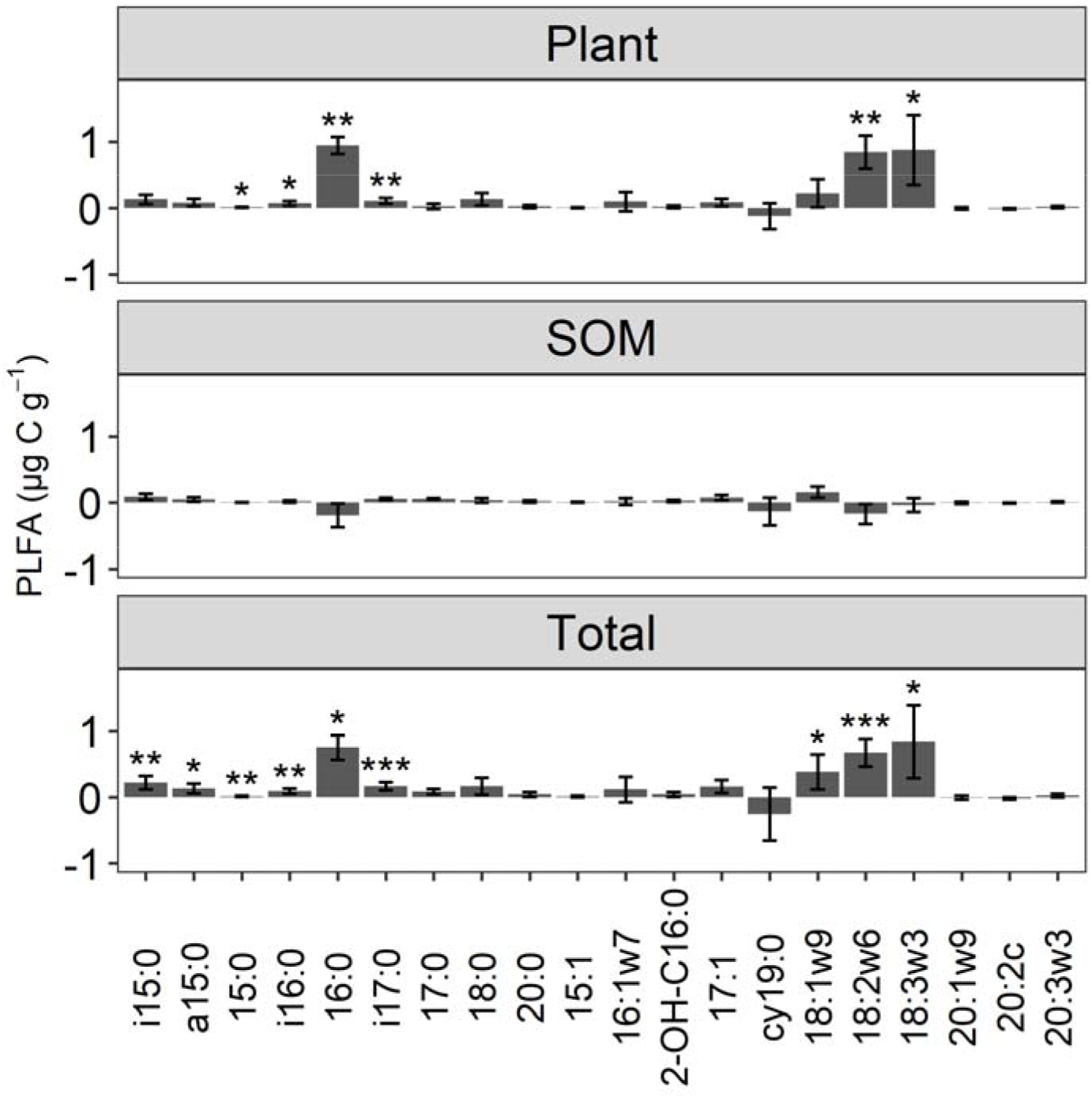
Differences in PLFA biomarker biomass between the mixture and the average of four non-mixture treatments. Positive values indicate that mixture has a greater biomass than the average of four non-mixtures. The three panels represent the crop residue derived PLFA biomass (Plant), the primed SOM derived PLFA biomass (SOM), and the total PLFA biomass (Total). Error bars are standard deviations. *, **, and *** indicates significantly different from zero at the level P < 0.05, 0.01, and 0.001, respectively.

One-way ANOSIM results demonstrated that microbial community structure in the mixture was significantly (*P* < 0.001, R = 0.326) different from that in the average of four non-mixture (Figure 4).

**Figure 4.**
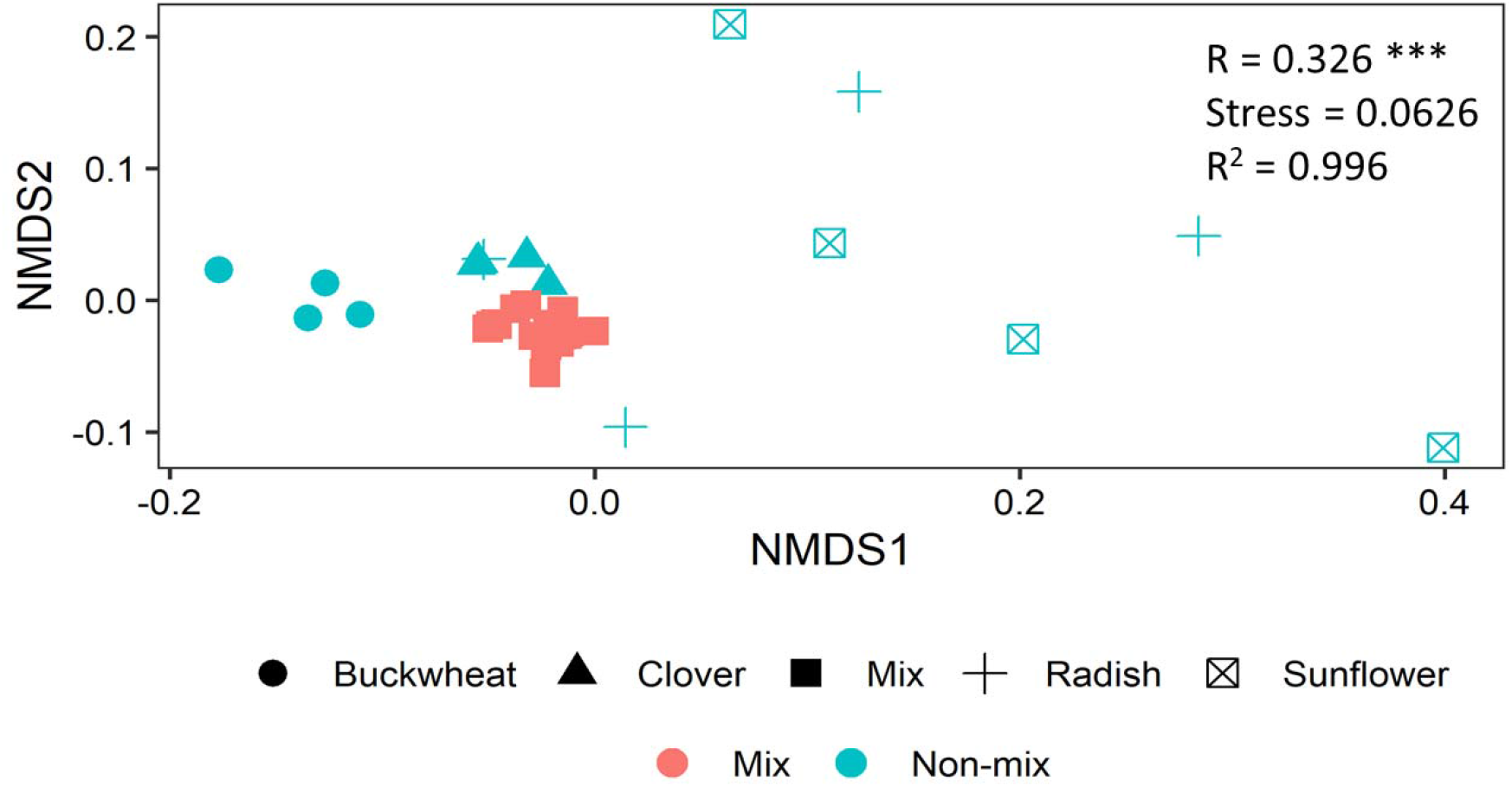
Microbial community structure in the mixture and non-mixture treatments. Non-metric multidimensional scaling (NMDS) on the Bray-Curtis distance on the Hellinger transformed PLFA data. Each symbol represents one sample. In the non-mixture treatment, different shapes represent different crop species. R value followed by “***” indicates significant (P < 0.001) difference between the mixture and the average of four non-mixture treatments analysed by one-way ANOSIM.

### 3.3 Mixing crop residues increased the PLFA biomass derived from crop residues

Applying ^13^C labelled cover residues allowed us to distinguish the microbial biomass derived from plant residues from other resources. The results showed that application of crop residues in the mixture and the average of four non-mixture treatments resulted in 6.34 and 2.73 µg C g^-1^ PLFA biomass derived from crop residues, respectively (Figure 1). We observed a significantly (*P* < 0.05) greater crop residue derived PLFA, by 132% (3.61 µg C g^-1^), in the mixture treatment, compared to the average of four non-mixture treatments (Figure 1).

The mixture exhibited a significantly (*P* < 0.01) greater biomass of general bacteria, Gram-positive bacteria, and fungi derived from crop residues, by 193% (1.15 µg C g^-1^), 86% (0.41 µg C g^-1^), and 158% (1.95 µg C g^-1^), respectively, compared to the average of four non-mixture treatment (Figure 2 and Table 2). The biomass of 16:0, 18:2ω6, and 18:3ω3 in the mixture treatment were significantly (*P* < 0.05) greater than those in the average of non-mixture treatments (Figure 3). Based on the differences in PLFA biomass derived from crop residues, there was no significant differences in either the fungi to bacteria ratio, or the Gram-positive to Gram-negative bacteria ratio, between mixture and the average of four non-mixture treatments (Figure S2 and S3).

### 3.4 Mixing crop residues did not increase the PLFA biomass derived from primed SOM

The PLFA biomass derived from primed SOM was the difference between total PLFA, PLFA in the control soil, and the PLFA derived from plant residues which was tracked by ^13^C. The results showed that the PLFA biomass derived from primed SOM was 3.90 and 3.82 µg C g^-1^ for the mixture and the average of four non-mixture treatments, respectively (Figure 1). Therefore, we observed only 2% (0.08 µg C g^-1^) greater PLFA biomass derived from primed SOM between the mixture and the average of four non-mixture treatments which was not statistically significant (Figure 1 and Table 2). None of the microbial groups or the corresponding PLFA biomarkers exhibited significant differences between the mixture and the average of four non-mixtures in terms of SOM priming (Figure 2 and 3). There were no significant differences in either the fungi to bacteria ratio or the Gram-positive to Gram-negative bacteria of PLFA derived from primed SOM between the mixture and the average of four non-mixture treatments (Figure S2 and S3).

## 4. Discussion

Following the addition of ^13^C-labelled residues, we compared the effect of mixing crop residues with the effect of applying the residues of a single crop species, on the soil microbial community composition and attributed the source of the C assimilated by the microbial biomass. Applying crop residues as a mixture resulted in a significantly (*P* < 0.05) greater microbial biomass compared to the average effect of applying each of the residues individually, indicating a synergistic effect of crop residue diversity on soil microbial biomass (Figure 1). This result was consistent with a microcosm experiment which found 38.2% higher soil microbial biomass N in soils receiving a mixture of residues than would be expected from residues of individual species (Mao et al., 2017). The mixture also exhibited greater total microbial biomass than any of the non-mixture treatments (Figure S4), suggesting its synergistic effect is more likely to be caused by facilitation between residues rather than by the disproportionate effect of individual species. The common explanation in the literature for why a mixture induces synergistic effect on decomposition rate is that N is transferred from low C/N residues to high C/N residues to satisfy microbial stochiometric requirement (Gartner and Cardon, 2004; Mao et al., 2017). If the availability of N is the limiting factor in our experiment, we would see the treatment receiving clover residues (which had the highest C/N ratio) exhibiting the smallest microbial biomass. On the contrary, the total microbial biomass in the treatment receiving clover residue (where C/N ratio was 32) was significantly (*P* < 0.05) greater than the treatment receiving buckwheat residue (where C/N ratio was 10) (Figure S4), implying that N transfer between high C/N and low C/N residues to satisfy microbial stoichiometric requirement may not be the reason for the observed synergistic effects of the mixture.

Although crop residues, such as buckwheat and sunflower in our study, contain high amount of N, they could also contain a considerable amount of plant secondary metabolites (e.g., tannins and terpenes) that suppress microbial resource assimilation (Gessner et al., 2010) or require specialized enzymes to be degraded (Chomel et al., 2016). When mixing residues, microorganisms could acquire energy via decomposing liable C from other residues to synthesize enzymes to degrade these secondary metabolites, and then anabolise subsequent monomers in their biomass (Wang et al., 2015). Additionally, micronutrients (e.g., Mg and Ca) were identified to play a paramount role in regulating nutrient transfer between residues (García-Palacios et al., 2016). This includes positive effects of Ca on the growth and activity of white rot fungi and significant effects of Mg on invertebrate (e.g., nematodes) diets which may influence the soil microbial community composition (García-Palacios et al., 2016). Thus, we suggested that the non-additive effect induced by the mixture was not predominantly controlled by residue bulk elemental composition (C/N ratio); instead, it could have been driven by the chemical composition of residues, including plant secondary metabolites and micronutrients.

We found that the greater microbial biomass in soils receiving residue mixtures, compared to individual residues, can largely be attributed to C assimilated directly from the plant residues, rather than C obtained by enhanced priming of SOM (Figure 1). Microbial communities preferentially mineralise labile compounds after crop residues are applied to soils to build their biomass rather than decomposing pre-existing SOM (Ball et al., 2014). Significantly greater crop residue-derived C was observed in biomarkers 16:0, 18:2ω6 and 18:3ω3 in the soils amended with crop residue mixtures, compared to soils receiving individual residues (Figure 3). This finding is supported by the knowledge that biomarker 16:0 represents microbial groups that primarily build their biomass from crop derived C (Wang et al., 2014). Although 16:0 was widely accepted as general bacterial biomarker, it also found in crops (Willers et al., 2015). There is a possibility that this biomarker was present as part of the tissues of our cover crops. Therefore, future studies combining stable isotope probing and DNA fingerprints would overcome this uncertainty associated with the PLFA approach (Blagodatskaya and Kuzyakov, 2013). We found that fungi (e.g., biomarker 18:2ω6 and 18:3ω3) were particularly efficient in assimilating a mixture of crop residue-derived C (Figure 3). This could be because fungal hyphae networks allow nutrients to be transported between microsites in the chemically and spatially heterogenous environment resulting from mixed residues (Ball et al., 2014). Furthermore, fungi can produce a wide range of extracellular enzymes that can degrade compounds which are recalcitrant for other microbes (Voriskova and Baldrian, 2013). By contrast, Gram-negative bacteria (e.g., 16:1ω5 and 16:1ω7) are good at incorporating SOM derived C into their biomass (Nottingham et al., 2009).

Although the application of crop residues did induce microbial assimilation of the native SOM, the magnitude of assimilation was not significantly changed by mixing crop residues (Figure 1 and 2). This observation was contrary to a previous assertion that mixtures of plant residues with a wide spectrum of labile compounds may support a higher microbial biomass and produce more extracellular enzymes, consequently enhancing the potential to prime the decomposition of recalcitrant compounds in SOM (Meier and Bowman, 2008). The lack of a marked mixture-induced priming effect could, however, be because the amount of C added was the same in all treatments. A recent meta-analysis, which analysed studies applying a range of organic C application rates, up to 3 mg C g^-1^, reported that the magnitude of the priming effect significantly increased with the increasing rate of additions, but was not affected by different residue types (Sun et al., 2019).

We made our observations only one day after crop residues were applied to soils because we were interested in identifying the microbial groups which incorporate plant derived C directly into their biomass by anabolism, rather than secondary turnover of C from microbial necromass, which could continue for months or years (Gunina and Kuzyakov, 2015). If incubating longer until the exhaustion of available C, then mixture treatments which have a larger microbial biomass, may facilitate a stronger priming effect because of the increased need for nutrients to maintain microbial survival (Yu et al., 2020).

Our study revealed that the incorporation of crop residue mixtures increased the total microbial biomass during the first day after application to a greater extent than would be expected by the addition of the same quantity of C from a single species residue. This additional microbial biomass is mostly derived from the crop residues themselves, rather than primed SOM. It is not clear whether the reason for this greater microbial biomass is due to faster metabolism of the residues, or higher carbon use efficiency (CUE) of the microorganisms responsible. We also found that soil respiration rates one day after applying crop residues were not significantly (*P* < 0.05) different between the mixture and the average of four non-mixtures (Figure S7). If the mechanism of higher CUE is responsible (or even partially responsible), then applying a mixture of crop residues to soils may help to increase SOC stocks in arable soils because more crop residue-derived C enters the microbial biomass and less is respired as CO_2_ (Figure S7) and the larger microbial biomass may result in a larger microbial necromass and, ultimately, more SOM. Therefore, land management practices that involve the incorporation of a mixture of crop residues (such as growing cover crop mixtures in polyculture after monoculture crop harvest and before planting the following monoculture crop) may be beneficial for increasing soil C stocks and climate change mitigation.

## 5. Conclusion

Mixing of crop residues produced a synergistic effect on total soil microbial biomass because fungi, general bacteria, and Gram-positive bacteria were able to incorporate plant-derived C directly into their biomass to a greater extent than when applying individual crop residues. These findings may be due to a better balance of nutrient inputs provided by mixtures, and broader niche for microorganism to colonise. Residue addition primed native SOM, but mixing residues resulted in an additive effect on microbial biomass C resulting from primed SOM. If the greater soil microbial biomass from residue mixtures results in greater microbial necromass and does not result in later priming of native SOM, then applying a mixture of crop residues to soils (e.g., incorporating or growing a cover crop polyculture) could be a sustainable agricultural practice to enhance SOC storage in arable soils.

## Acknowledgements

This research was supported by a Biotechnology and Biological Sciences Research Council New Investigator Grant (BB/R006989/1) awarded to Tom Sizmur. The authors wish to thank Paul Brown from Kings Crops, a division of Frontier Agriculture Ltd, for supplying cover crop seed, and the Isolife (Wageningen, Netherland) for growing and labelling the cover crops. We acknowledge the assistance of Andrew Dodson, Anne Dudley, Karen Gutteridge, Fengjuan Xiao, Marta O’Brian, Sean Coole with laboratory analysis, and Dedy Antony, Alex Adetunji Adekanmbi and Chinonso Chukwuma Ogbuagu with soil sampling.

## Supplementary information

A Supplementary information document is provided which contains 3 tables and 7 figures

