## Supplementary information for "Mixing crop residues induces a synergistic effect on microbial biomass and an additive effect on soil organic matter priming"

### This supplementary information contains:

**3 Tables (Table S1, S2 and S3)**

**7 Figures (Figure S1, S2, S3, S4, S5, S6 and S7)**

**Table S1** The chemical composition of $^{13}C$ labelled and unlabelled crop residues. The $^{13}C$ atom percent in the unlabelled crops was not measured.

| Crop | Family | C (%) | N (%) | C/N ratio | $^{13}C$ atom percent (%) |
|---|---|---|---|---|---|
| **$^{13}C$ labelled plants** | | | | | |
| Buckwheat | Polygonaceae | 38.77 | 2.73 | 14 | 6.7 |
| Clover | Fabaceae | 40.70 | 1.00 | 41 | 7.8 |
| Radish | Brassicaceae | 35.96 | 2.73 | 13 | 7.8 |
| Sunflower | Asteraceae | 37.46 | 2.65 | 14 | 8.0 |
| **Unlabelled plants** | | | | | |
| Buckwheat | Polygonaceae | 37.12 | 4.28 | 9 | \ |
| Clover | Fabaceae | 39.30 | 1.28 | 31 | \ |
| Radish | Brassicaceae | 37.19 | 1.76 | 21 | \ |
| Sunflower | Asteraceae | 38.48 | 1.66 | 23 | \ |

**Table S2** The attribution of PLFA biomarkers to microbial groups

| Microbial groups | PLFA biomarkers | References |
|---|---|---|
| Gram-positive (G+) bacteria | i15:0, a15:0, i16:0, i17:0 | (Mbuthia et al., 2015; Wilkinson et al., 2002; Zelles, 1999; Zheng et al., 2018) |
| Gram-negative (G-) bacteria | 15:1, 16:1 ω7, 17:1, cy19:0, 2-OH-C16:0 | |
| Fungi | 18:2 ω6, 18:1 ω9, 18:3ω3, 20:1 ω9c | |
| Protozoa | 20:2c, 20:3ω3 | |
| General bacteria | 15:0, 16:0, 17:0, 18:0, 20:0 | |

**Table S3 Total microbial biomass (µg C $g^{-1}$) in all treatments measured by GC-FID. Total is the sum of all the four microbial groups. G+ and G- are Gram-positive and Gram-negative bacteria. F/B ratio and G+/G- ratio represent the fungi to bacteria ratio, and the Gram-positive to Gram-negative bacteria ratio. Mean ± standard deviation (n = 4). *, **, and *** mean significant difference from the control soil at the level $P$ < 0.05, 0.01 and 0.001, respectively. Refer to Table 1 for treatment codes.**

| Treatment | General bacteria | G+ bacteria | G- bacteria | Fungi | Protozoa | Total | F/B ratio | G+/G- ratio |
|---|---|---|---|---|---|---|---|---|
| MB | 4.13±0.59*** | 3.73±0.51*** | 2.20±0.36 | 6.26±0.86*** | 0.08±0.02 | 16.40±2.32*** | 0.62±0.02 | 1.70±0.06 |
| MC | 4.69±0.55*** | 4.09±0.41*** | 2.25±0.29* | 7.30±0.82*** | 0.09±0.01 | 18.42±1.77*** | 0.66±0.03* | 1.84±0.32 |
| MR | 4.76±0.48*** | 4.26±0.52*** | 2.43±0.31** | 7.38±0.73*** | 0.09±0.01 | 18.93±1.99*** | 0.65±0.03* | 1.75±0.03 |
| MS | 4.33±0.32*** | 3.90±0.30*** | 2.23±0.16* | 6.67±0.41*** | 0.09±0.01 | 17.22±1.14*** | 0.64±0.02* | 1.75±0.01 |
| NB | 3.06±0.19 | 2.79±0.16 | 1.41±0.10 | 4.99±0.59** | 0.03±0.04 | 12.29±1.04* | 0.69±0.05** | 1.99±0.05 |
| NC | 3.51±0.32** | 3.53±0.31*** | 1.89±0.17 | 5.12±0.30** | 0.08±0.01 | 14.14±1.09*** | 0.57±0.02 | 1.86±0.01 |
| NR | 4.02±0.61*** | 3.94±0.53*** | 2.77±0.76*** | 6.04±2.18*** | 0.10±0.12 | 16.88±3.35*** | 0.57±0.18 | 1.48±0.31 |
| NS | 3.05±1.20 | 3.22±0.24* | 2.68±1.16** | 3.86±0.82 | 0.08±0.04 | 12.89±0.57** | 0.44±0.12 | 1.40±0.60 |
| PB | 3.11±0.48 | 2.64±0.41 | 1.42±0.21 | 4.57±0.50* | 0.03±0.04 | 11.77±1.48* | 0.64±0.10* | 1.86±0.02 |
| PC | 3.02±0.27 | 2.99±0.13 | 1.54±0.06 | 4.51±0.27* | 0.02±0.03 | 12.08±0.71* | 0.60±0.01 | 1.94±0.03 |
| PR | 3.69±0.41*** | 3.69±0.33*** | 2.06±0.21 | 5.36±0.40** | 0.05±0.03 | 14.84±1.39*** | 0.57±0.02 | 1.79±0.02 |
| PS | 2.96±0.36 | 2.74±0.35 | 1.46±0.17 | 4.48±0.60* | 0.03±0.04 | 11.66±1.49* | 0.62±0.02 | 1.88±0.05 |
| Soil | 1.80±0.27 | 2.20±0.27 | 1.15±0.14 | 2.36±0.32 | 0.00±0.00 | 7.50±0.99 | 0.46±0.01 | 1.92±0.03 |

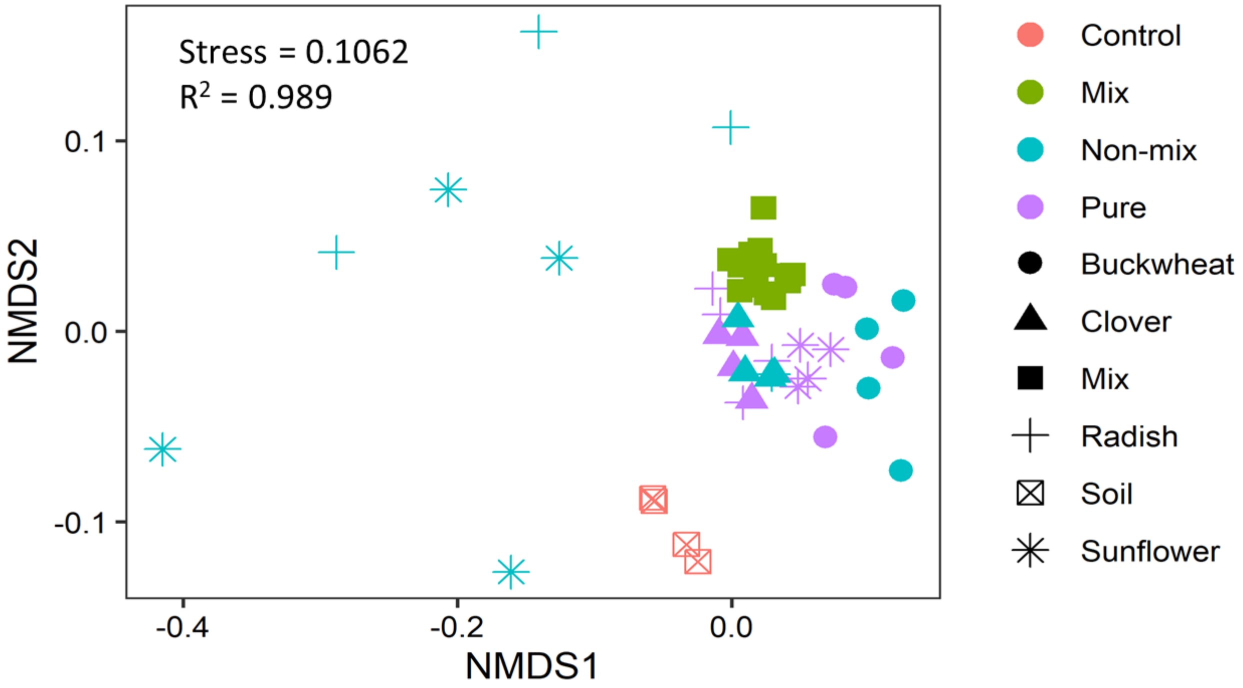


**Figure S1** Bacterial community structure. Non-metric multidimensional scaling (NMDS) on the Bray-Curtis distance on the Hellinger transformed PLFA data. Dots the NMDS scores of samples. For the pure and non-mixture treatment, different shape represents the plant species in that treatment. One-way ANOSIM indicates a significant (R = 0.899, $P < 0.001$) difference between all the pure and control treatments. In the pure treatments, different plant species induced significant ($P < 0.05$) different microbial community composition.

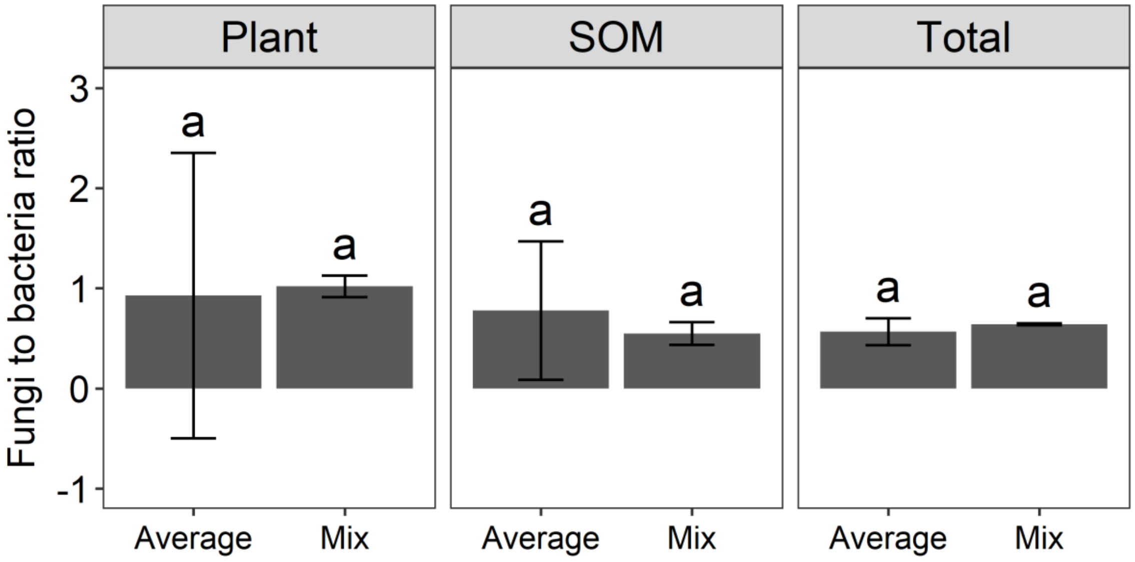


**Figure S2** The ratio of fungi to bacteria biomass based on the PLFA derived from crop residues (Plant), primed soil organic matter (SOM), and total PLFA (Total). Average is the average of four non-mixture treatments. Different letters within the same panel mean significant difference between mixture and the average of four non-mixture treatments at $P < 0.05$. The error bars are standard deviations (n = 4).

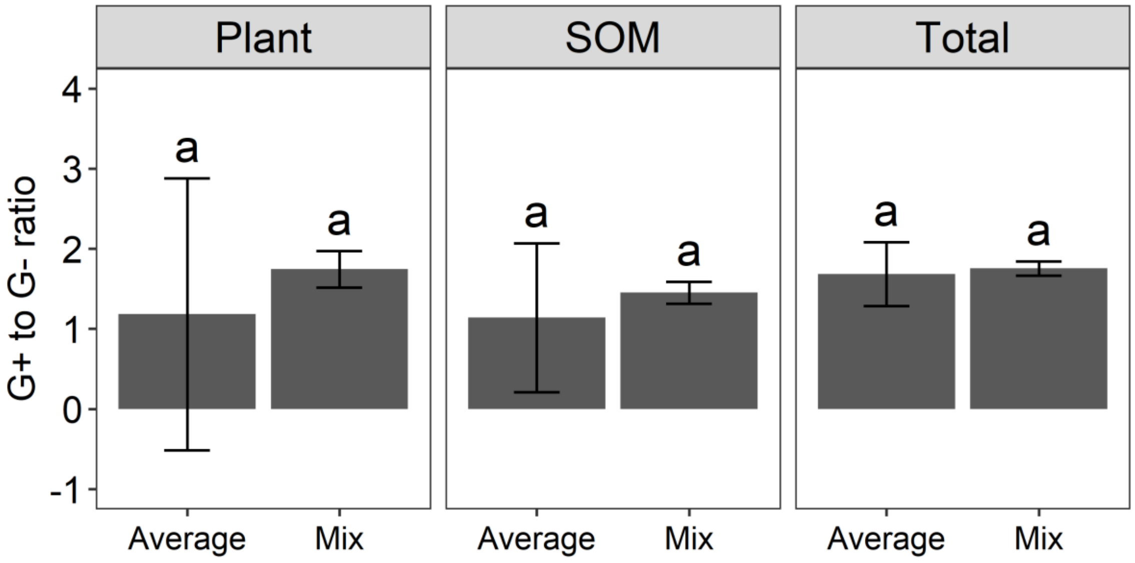


**Figure S3** The ratio of Gram-positive bacteria (G+) to Gram-negative bacteria (G-) biomass based on the PLFA derived from crop residues (Plant), primed soil organic matter (SOM), and total PLFA (Total). Average is the average of four non-mixture treatments. Different letters within the same panel mean significant difference between mixture and the average of four non-mixture treatments at $P < 0.05$. The error bars are standard deviations (n = 4).

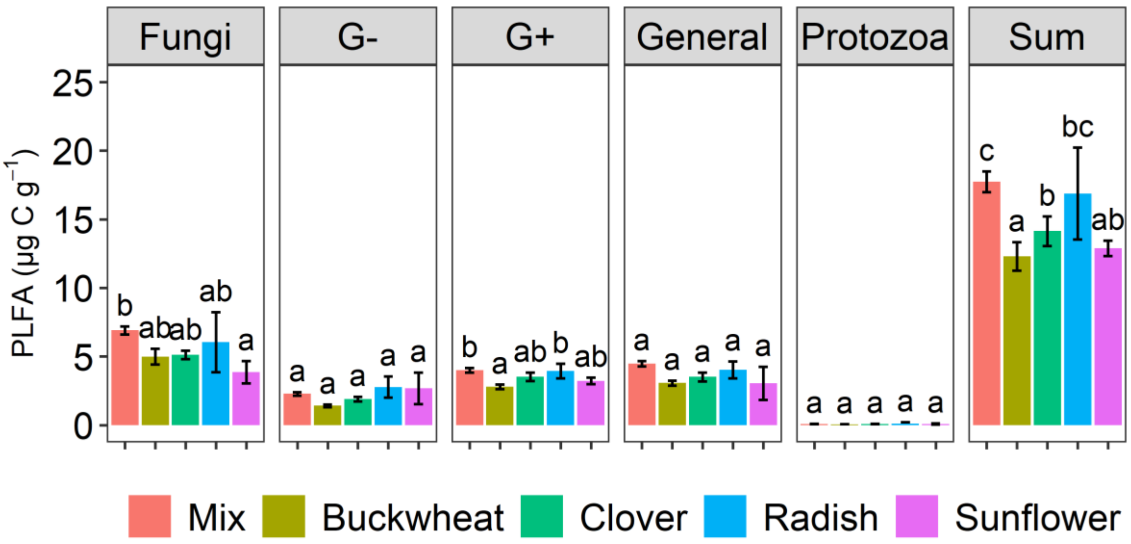


**Figure S4** Total PLFA biomass in the mixture, and non-mixture (buckwheat, clover, radish, and sunflower) treatments. Sum is the sum of microbial biomass including all the microbial groups. G+, G-, and General represent Gram-positive, Gram-negative, and general bacteria. Error bars are standard deviations (n = 4). Different letters above bars indicate significant differences in PLFA biomass between treatments for that microbial group at *P* < 0.05.

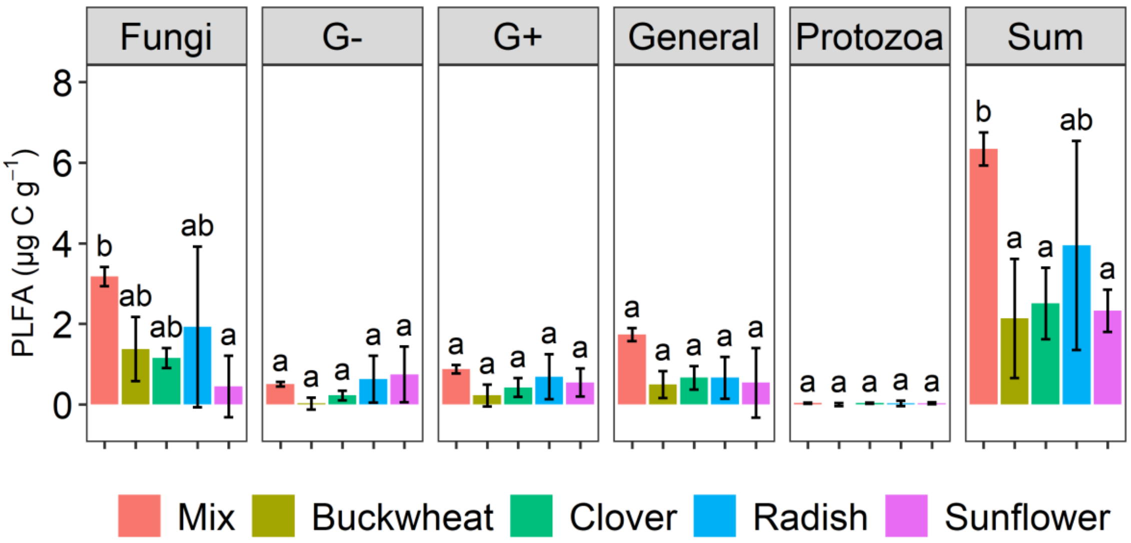


**Figure S5** Crop residue- derived PLFA biomass in the mixture, and non-mixture (buckwheat, clover, radish, and sunflower) treatments. Sum is the sum of microbial biomass including all the microbial groups. G+, G-, and General represent Gram-positive, Gram-negative, and general bacteria. Error bars are standard deviations (n = 4). Different letters above bars indicate significant differences in PLFA biomass between treatments for that microbial group at *P* < 0.05.

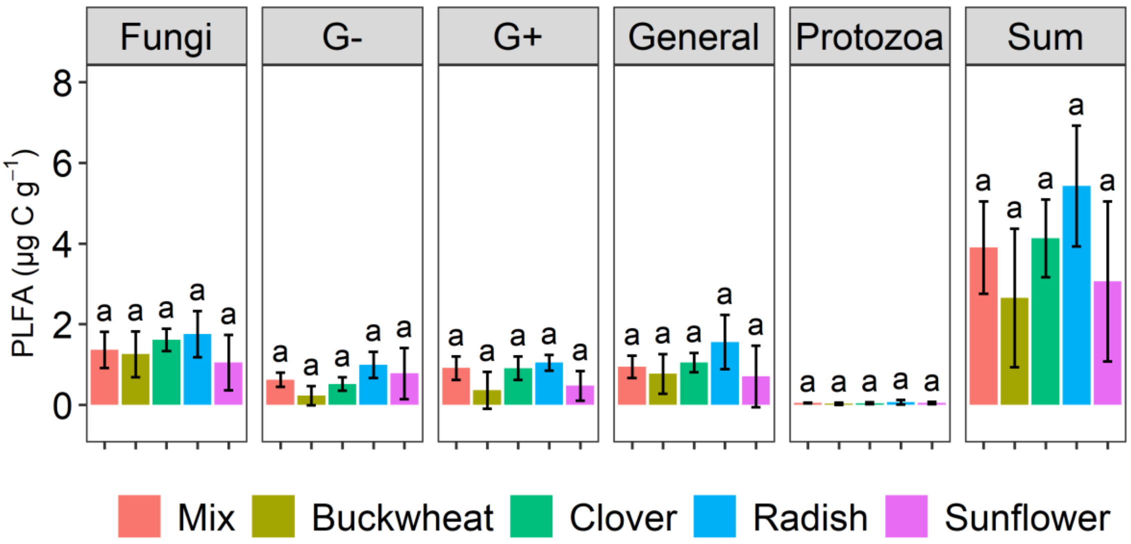


**Figure S6** SOM (soil organic matter) derived PLFA biomass in the mixture, and non-mixture (buckwheat, clover, radish, and sunflower) treatments. Sum is the sum of microbial biomass including all the microbial groups. G+, G-, and General represent Gram-positive, Gram-negative, and general bacteria. Error bars are standard deviations (n = 4). Different letters above bars indicate significant differences in PLFA biomass between treatments for that microbial group at ***P*** < 0.05.

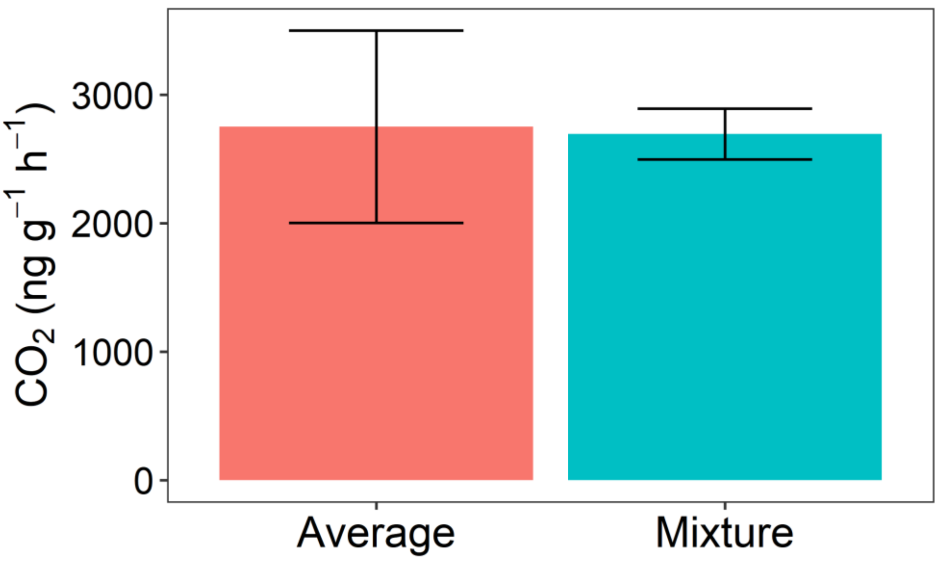


**Figure S7** Respiration rate in the mixture and the average of four non-mixture treatments one day after applying crop residues. The respiration rates were not significant different between the mixture and the average of four non-mixtures. Error bars are standard deviations.
